# An optimality principle for locomotor central pattern generators

**DOI:** 10.1101/2019.12.30.890152

**Authors:** Hansol X. Ryu, Arthur D. Kuo

**Author notes:** Corresponding author: H. X. Ryu. Co-corresponding author: A. D. Kuo.

## Abstract

Two types of neural circuits contribute to legged locomotion: *central pattern generators* (CPGs) that produce rhythmic motor commands (even in the absence of feedback, termed “fictive locomotion”), and *reflex circuits* driven by sensory feedback. Each circuit alone serves a clear purpose, and the two together are understood to cooperate during normal locomotion. The difficulty is in explaining their relative balance objectively within a control model, as there are infinite combinations that could produce the same nominal motor pattern. Here we propose that optimization in the presence of uncertainty can explain how the circuits should best be combined for locomotion. The key is to re- interpret the CPG in the context of state estimator-based control: an internal model of the limbs that predicts their state, using sensory feedback to optimally balance competing effects of environmental and sensory uncertainties. We demonstrate use of optimally predicted state to drive a simple model of bipedal, dynamic walking, which thus yields minimal energetic cost of transport and best stability. The internal model may be implemented with classic neural half-center circuitry, except with neural parameters determined by optimal estimation principles. Fictive locomotion also emerges, but as a side effect of estimator dynamics rather than an explicit internal rhythm. Uncertainty could be key to shaping CPG behavior and governing optimal use of feedback.

**New and Noteworthy:** Sensory feedback modulates the central pattern generator (CPG) rhythm in locomotion, but there lacks an explanation for how much feedback is appropriate. We propose destabilizing noise as a determinant, where an uncertain environment demands more feedback, but noisy sensors demand less. We reinterpret the CPG as an internal model for predicting body state despite noise. Optimizing its feedback yields robust and economical gait in a walking model, and explains the advantages of feedback-driven CPG control.

## Introduction

A combination of two types of neural circuitry appears responsible for the basic locomotory motor pattern. One type is the central pattern generator (CPG; Figure 1A), which generates pre-programmed, rhythmically timed, motor commands (1–3). The other is the reflex circuit, which produces motor patterns triggered by sensory feedback (Figure 1C). Although they normally work together, each is also capable of independent action. The intrinsic CPG rhythm patterns can be sustained with no sensory feedback and only a tonic, descending input, as demonstrated by observations of fictive locomotion (4,5). Reflex loops alone also appear capable of controlling locomotion (1), particularly with a hierarchy of loops integrating multiple sensory modalities for complex behaviors such as stepping and standing control (6,7). We refer to the independent extremes as pure feedforward control and pure feedback control. Of course, within the intact animal, both types of circuitry work together for normal locomotion (Figure 1B; (8)). But this cooperation also presents a dilemma, of how authority should optimally be shared between the two (9).

**Figure 1.**
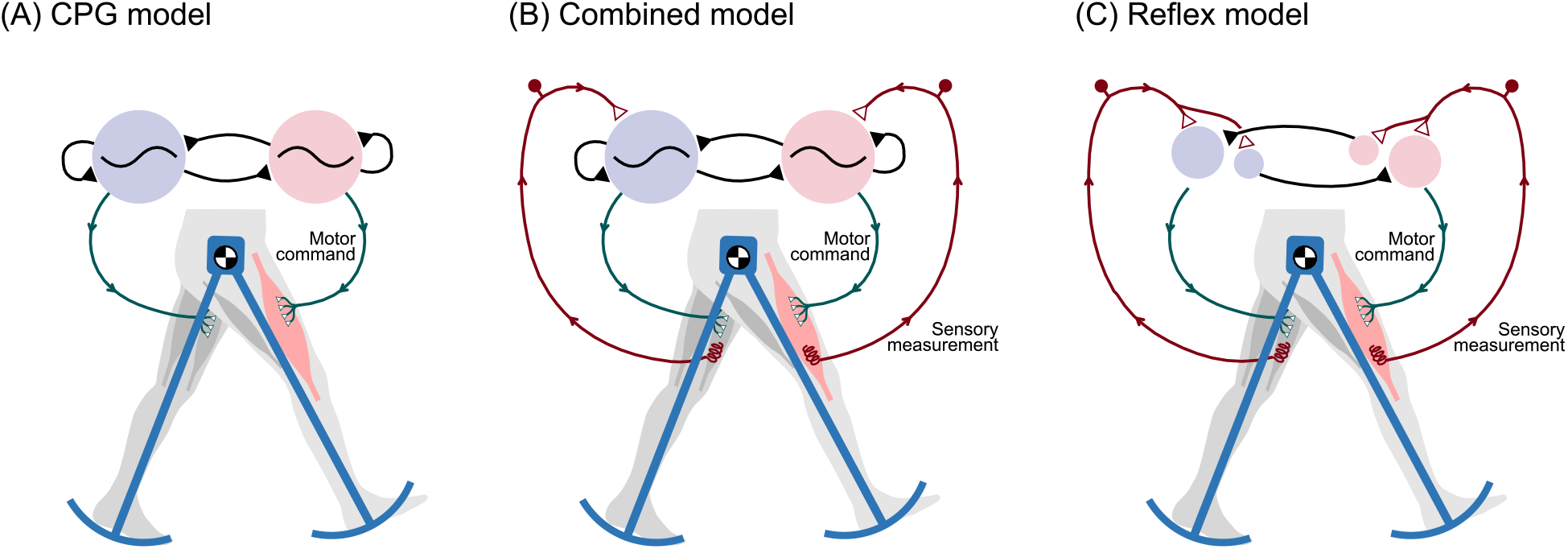
Three ways to control bipedal walking. (A) The central pattern generator (CPG) comprises neural oscillators that can produce rhythmic motor commands, even in the absence of sensory feedback. Rhythm can be produced by mutually inhibiting neural half-center oscillators (shaded circles). (B) In normal animal locomotion, the CPG is thought to combine an intrinsic rhythm with sensory feedback, so that the periphery can influence the motor rhythm. (C) In principle, sensory feedback can also control and stabilize locomotion through reflexes, without need for neural oscillators. The extreme of (A) CPG control without feedback is referred to here as pure feedforward control, and the opposite extreme (C) with no oscillators as pure feedback control. Any of these schemes could potentially produce the same nominal locomotion pattern, but some (B) combination of feedforward and feedback appears advantageous.

The combination of central pattern generators with sensory feedback has been explored in computational models. For example, some models have added feedback (10–13) to biologically-inspired neural oscillators (e.g., 14), which employ networks of mutually inhibiting neurons to intrinsically produce alternating bursts of activity. Sensory input to the neurons can change network behavior based on system state, such as foot contact and limb or body orientation, to help respond to disturbances. The gain or weight of sensory input determines whether it slowly entrains the CPG (15), or whether it resets the phase entirely (16,17). Controllers of this type have demonstrated legged locomotion in bipedal (18) and quadrupedal robots (19,20), and even swimming and other behaviors in others (21). A general observation is that feedback improves robustness, such as ability to traverse different terrains (22). Although such CPG model can certainly be devised for stable locomotion, the amount of feedback is often tuned ad hoc to reproduce biological behavior. Most CPG models to date lack an objective way to determine how much feedback to use, and may thus be suboptimal.

Optimization principles offer a means for a model to be uniquely defined by quantitative and objective performance measures (23). Thus far, human-like models that learn or optimize control generally favor state-based control (24–26), meaning that the control command is a function of system state (e.g., positions and velocities of limbs). In fact, reinforcement learning and other optimization approaches (e.g., dynamic programming (27,28)) are typically expressed solely in terms of state, and do not even have provision for time as an explicit input. They have no need for, nor even benefit from, an internally generated rhythm. But feedforward is clearly important in biological CPGs, suggesting that some insight is missing from these optimal control models.

There may be a principled reason for a biological controller not to rely on state, as measured, alone. Realistically, a system’s state can only be known imperfectly, due to noisy and imperfect sensors. The solution is state estimation (29), in which an internal model of the body is used to predict the expected state and sensory information, and feedback from actual sensors is used to correct the state estimate.

Actual robots (e.g., bipedal Atlas (30) and quadrupedal BigDog (31)) gain high performance and robustness through state estimation driving state-(estimator)-based control. In fact, state-estimator control may be optimized for a noisy environment (29), and has been proposed as a model for biological systems (32–34). The internal model for state estimation is not usually regarded as relevant to the biological CPG’s feedforward, internal rhythm. But we have proposed that it can potentially produce a CPG-like rhythm under conditions simulating fictive locomotion (35). Here the rhythmic output is interpreted not as the motor command *per se*, but as a state estimate that drives the motor command. We demonstrated this concept with a simple model of rhythmic leg motions (35) and a preliminary walking model (36). This suggests that an walking model designed objectively with state estimator-based control might produce CPG-like rhythms that are entirely compatible with optimal performance.

The purpose of the present study was to test an estimator-based CPG controller with a dynamic walking model. We devised a simple state-based control scheme, producing stance and swing leg torques as a function of the leg states. Assuming a noisy system, we devised a state estimator for leg states, with a discrete switch between stance and swing dynamics. The combination of control and estimation thus define our interpretation of a CPG controller that incorporates sensory feedback. In fact, this same controller may be realized in the form of a biologically-inspired half center oscillator (14), with similar neuron-like dynamics. We expected that minimum state estimation error would allow this model to walk with optimal performance, in terms of measures such as mechanical cost of transport. Scaling the sensory feedback either higher or lower than optimal would be expected to yield poorer performance. Such a model may conceptually explain how CPGs could optimally incorporate sensory feedback.

## Results

### Central pattern generator controls a dynamic walking model

The CPG controller produced a periodic gait with a model of human-like dynamic walking (Figure 2A). The model was inspired by the mechanics and energetics of humans (37), whose legs have pendulum-like passive dynamics (38). Acting on the legs were torque commands (*T*_1_ and *T*_2_, Figure 2B) from the CPG, designed to yield a periodic gait (Figure 2C), by restoring energy dissipated with each step’s ground contact collision (39,40). The leg angles and the ground contact condition (“GC”, 1 for contact, 0 otherwise; Figure 2C) were treated as measurements to be fed back to the CPG. Each leg’s states (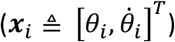) described a periodic orbit or limit cycle (Figure 2D), which was locally stable for zero or mild disturbances, but could easily be perturbed enough to make it fall (Figure 2E).

**Figure 2.**
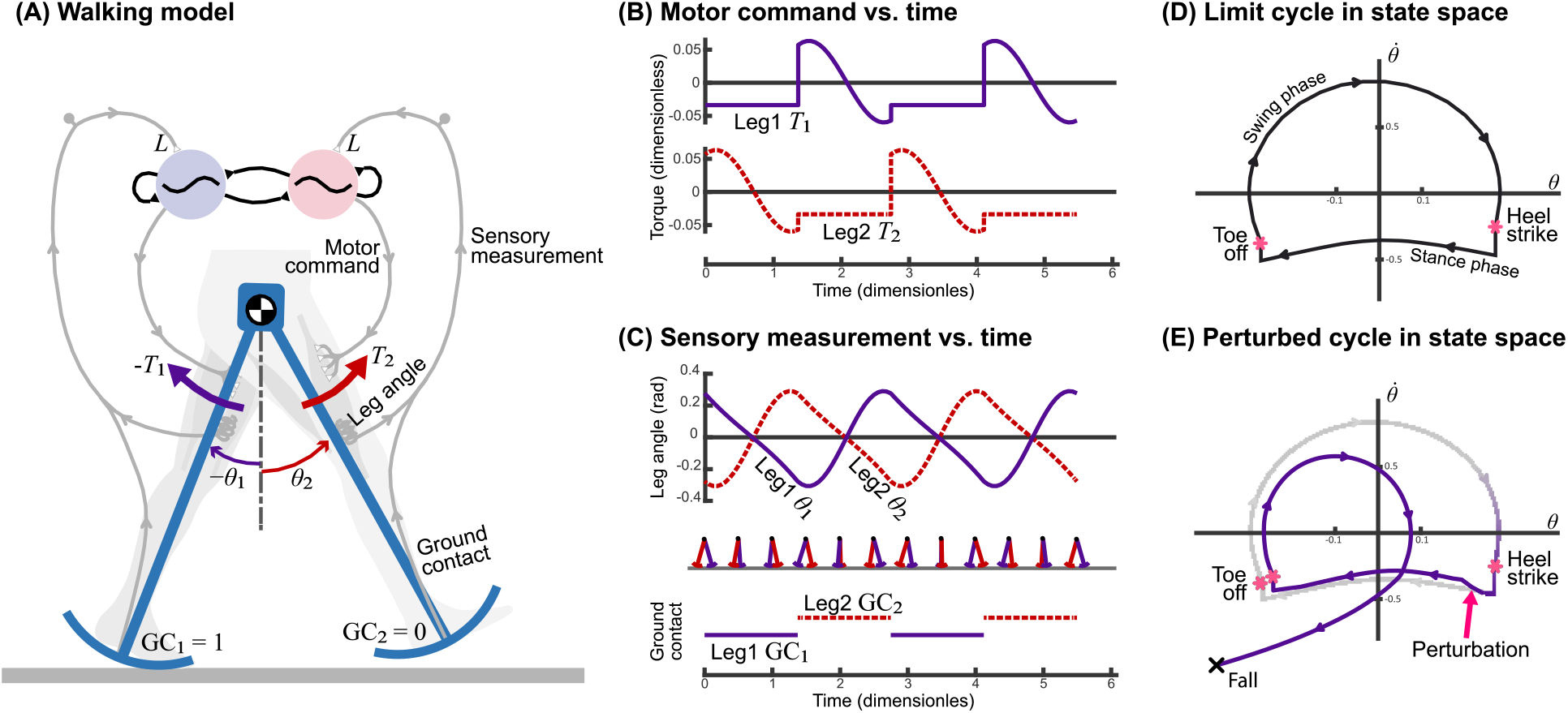
Dynamic walking model controlled by CPG controller with feedback. (A) Pendulum-like legs are controlled by motor commands for hip torques *T*_1_ and *T*_2_, with sensory feedback of leg angle and ground contact “GC” relayed back to controller. (B) Controller produces alternating motor commands vs. time, which drive (C) leg movement *θ*. Sensory measurements of leg angle and ground contact in turn drive the CPG. (D) Resulting motion is a nominal periodic gait (termed a “limit cycle”) plotted in state space 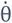 vs *θ*. (E) Discrete perturbation to the limit cycle can cause model to fall.

The resulting gait had approximately human-like parameters when walking without noisy disturbances. The nominal walking speed was equivalent to 1.25 m/s with step length 0.55 m (or normalized 0.4 (*gl*)^0.5^ and 0.55 *l*, respectively; *g* is gravitational constant, *l* is leg length). The corresponding mechanical cost of transport was 0.053, comparable to other passive and active dynamic walking models (e.g., (41–43)).

This controller had three important features for the analyses that follow. First, the gait had dynamic, pendulum-like leg behavior similar to humans (37,38). Second, the controller stabilized walking, meaning ability to withstand minor perturbations due to its state-based control. And third, the overall amount of feedback could be varied continuously between either extreme of pure feedforward and pure feedback (sensory gain *L* ranging zero to infinity), while always producing the same nominal gait under noiseless conditions. This was to facilitate study of how parametric variation of sensory feedback affects performance, particularly under noisy conditions.

### Pure feedforward and pure feedback both susceptible to noise

The critical importance of sensory feedback was demonstrated with a disturbance acting on the legs (Figure 3A). Termed *process noise*, it represents not only disturbances but also any uncertainty in the environment or internal model. For this demonstration, the disturbance consisted of a single impulsive hip torque acting on a leg. The pure feedforward controller failed to recover (Figure 3A left), and would fall within about two steps. Its perturbed leg and ground contact states became mismatched to the nominal rhythm, which in pure feedforward does not respond to state deviations. In contrast, the feedback controller could recover from the same perturbation (Figure 3A right) and return to the nominal gait. Feedback control is driven by system state, and therefore automatically alters the motor command in response to perturbations. Our expectation is that even if a feedforward control is stable under nominal conditions (with zero or mild disturbances), a feedback controller could generally be designed to be more robust.

**Figure 3.**
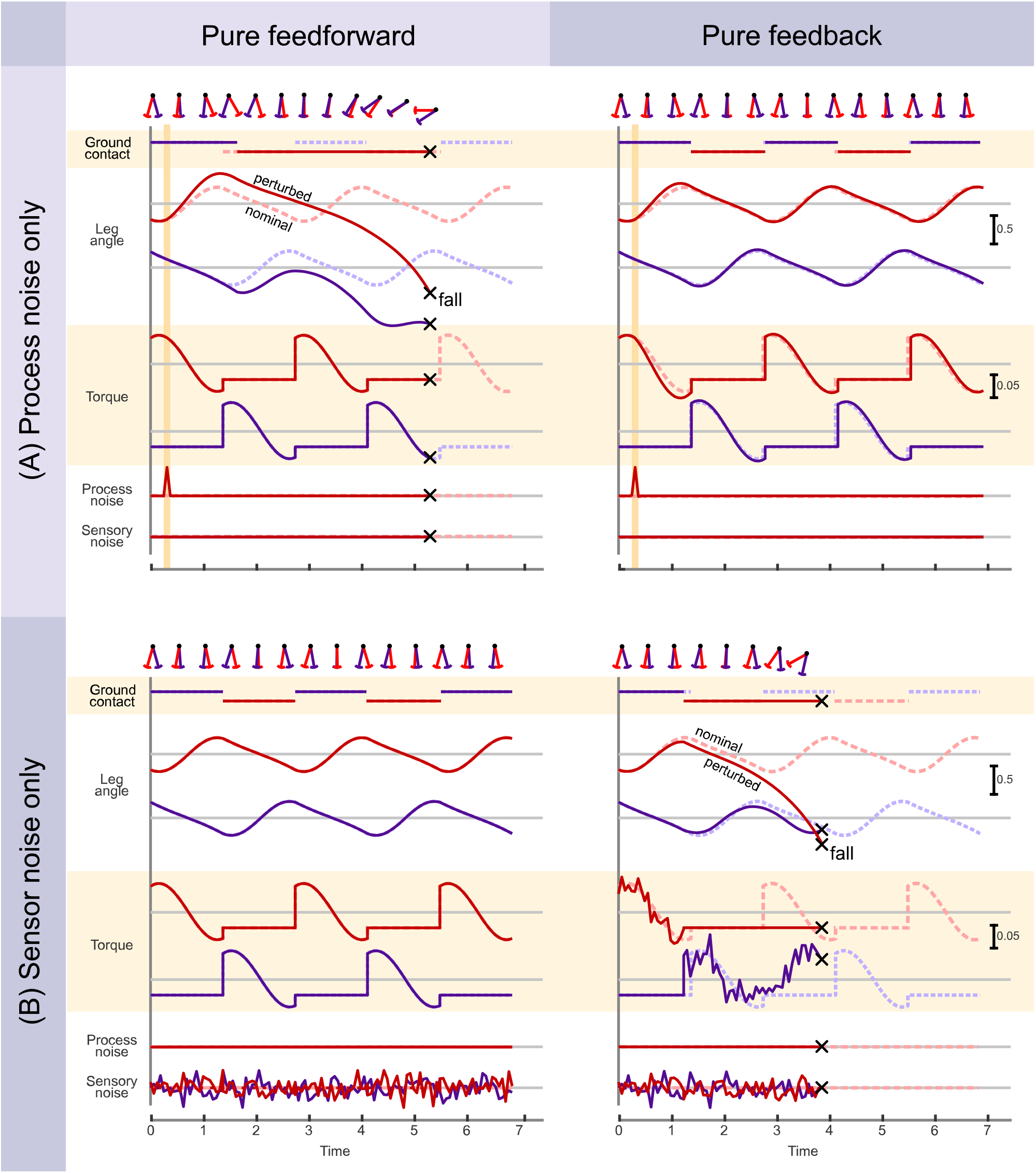
Effect of (A) process and (B) sensor noise on pure feedforward and pure feedback conditions (left and right columns, respectively). Plots show ground contact condition, leg angles, commanded leg torques, and noise levels vs. time, including both the nominal condition without noise (dashed lines), and the perturbed condition with noise (solid lines). With an impulsive, process noise disturbance, pure feedforward control tended to fall, whereas pure feedback was quite stable. With sensor noise alone, pure feedforward was unaffected, but pure feedback tended to fall.

We also applied an analogous demonstration with sensor noise (Figure 3B). Adding continuous *sensor noise* to sensory measurements had no effect on pure feedforward control (Figure 3B left), which ignores sensory signals entirely. But pure feedback was found to be sensitive to noise-corrupted measurements, and would fall within a few steps (Figure 3B right). This is because erroneous feedback would trigger erroneous motor commands not in accordance with actual limb state. The combined result was that both pure feedforward and pure feedback control had complementary weaknesses. They performed identically without noise, but each was unable to compensate for its particular weakness, either process noise or sensor noise. Feedback control can be robust, but it needs accurate state information.

### Equivalence between neural half-center oscillators and a state estimator

We determined two apparently different representations for the same CPG model. This first was a biologically inspired, neural oscillator (Figure 4A) representation, with two mutually inhibiting half-center oscillators, one driving each leg (*i* = 1 for left leg, *i* = 2 for right leg). Each half-center had a total of three neurons, one a primary neuron with standard second-order dynamics (states *u* and *v*). Its output drove the second neuron (*α*) producing the motor command to the ipsilateral leg. The third neuron was responsible for relaying ground contact (“*c*”) sensory information, to both excite the ipsilateral primary neuron and inhibit the contralateral one. This model was intended to resemble previous CPG models (10–13) demonstrating incorporation of sensory feedback into an intrinsic rhythm.

**Figure 4.**
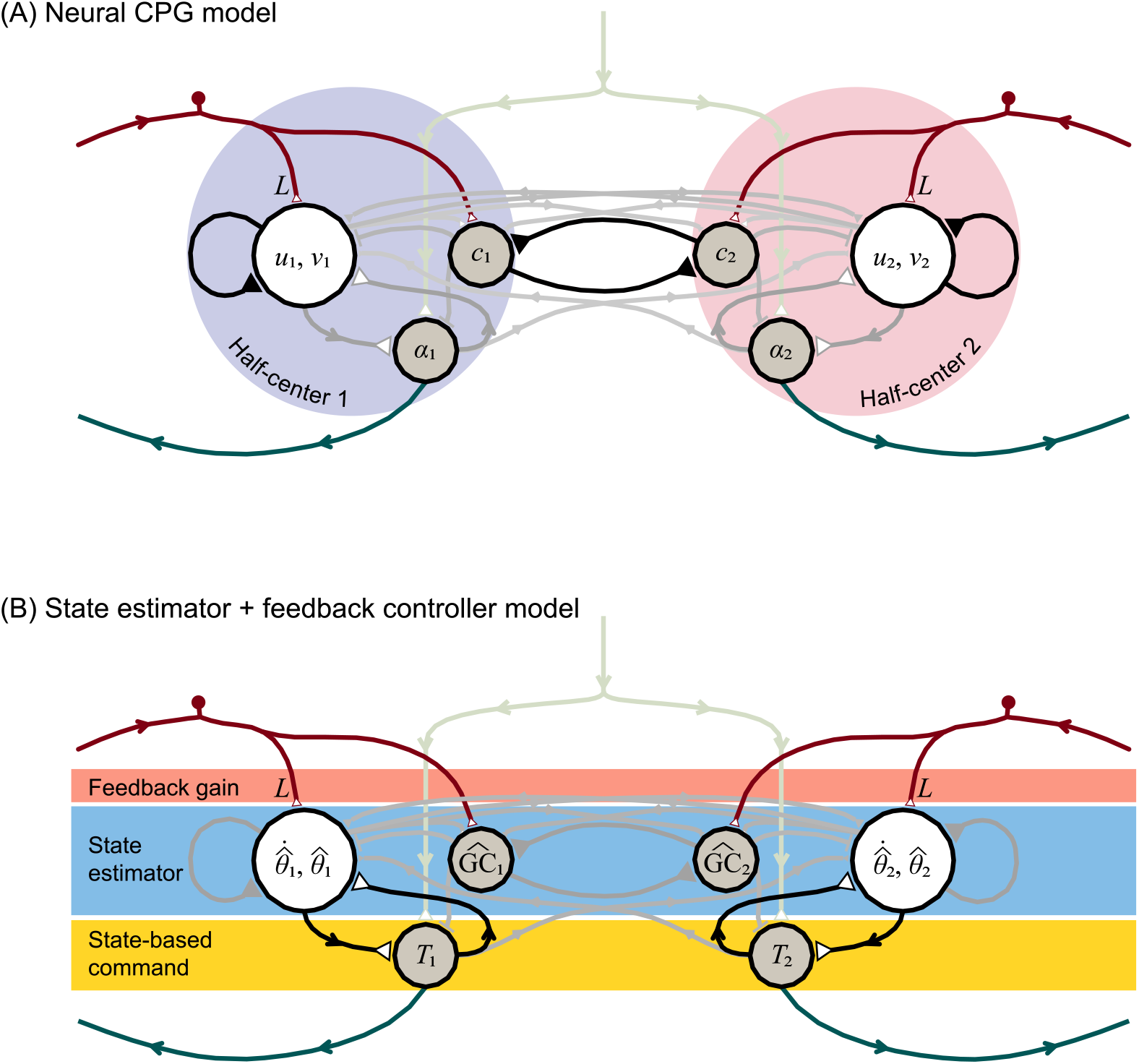
Locomotion control circuit interpreted in two representations: (A) Neural central pattern generator with mutually inhibiting half-center oscillators, and as (B) state estimator with feedback control. Each half-center has a primary neuron with two states (*u* and *v*, respectively), an auxiliary neuron *c* for registering ground contact, and an alpha motoneuron *α* driving leg torque commands. Inputs include a tonic descending drive, and afferent sensory data with gain *L*. State estimator acts as second-order internal model of leg dynamics to estimate leg states 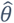 (hat symbol denotes estimate) and ground contact 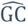, which drive state-based command *T*. All estimator parameters including sensory feedback *L* are designed for minimum mean-square estimation error. Leg dynamics have nonlinear terms (see Methods) of small magnitude (thin grayed lines).

The same CPG architecture was then re-interpreted in a second, control systems framework (Figure 4B), while changing none of the neural circuitry. Here, the structure was not treated as half-center oscillators, but rather as three neural stages from afferent to efferent. The first stage receiving sensory feedback signal was interpreted as a feedback gain *L* (upper rectangular block, Figure 4B), modulating the behavior of the second stage, interpreted as a state estimator (middle rectangular block, Figure 4B) acting as an internal model of leg dynamics. Its output was interpreted as the state estimate, which was fed into the third, state-based motor command stage (lower rectangular block, Figure 4B). In this interpretation, the three stages correspond with a standard control systems architecture for a state estimator driving state feedback control. In fact, the neural connection weights of the half-center oscillators were determined by, and are therefore specifically equivalent to, a state estimator driving motor commands to the legs.

The two representations provide complementary insights. The half-center model shows how sensory information can be incorporated into and modulate a CPG rhythm. Although the model can be tuned for a desired behavior, there is generally no objective and unique means to determine the neural weights. The state estimator-based model offers a means to determine the neural weights and parameters for the best performance, without need for ad hoc parameter tuning. This half-center model with feedback could thus be regarded as optimal.

### Sensory feedback gain *L* optimized by state estimation

We next examined walking performance in the presence of both process and sensory noise, while varying sensory gain *L* above and below optimal (Figure 5). We intentionally applied a substantial amount of noise (with fixed covariances), sufficient to topple the model. This was to demonstrate how walking performance can be improved with appropriate sensory gain *L*, unlike the noiseless case where the model always walks perfectly. With noise, both pure feedforward and pure feedback control yielded poor performance, as quantified by mechanical cost of transport, step variability, mean time between falls, and state estimator error (Figure 5). Better performance was achieved by varying sensory feedback *L* continuously between these extremes. The combination of feedforward and feedback, where the CPG rhythm was modulated by sensory information, performed better than either extreme alone.

**Figure 5.**
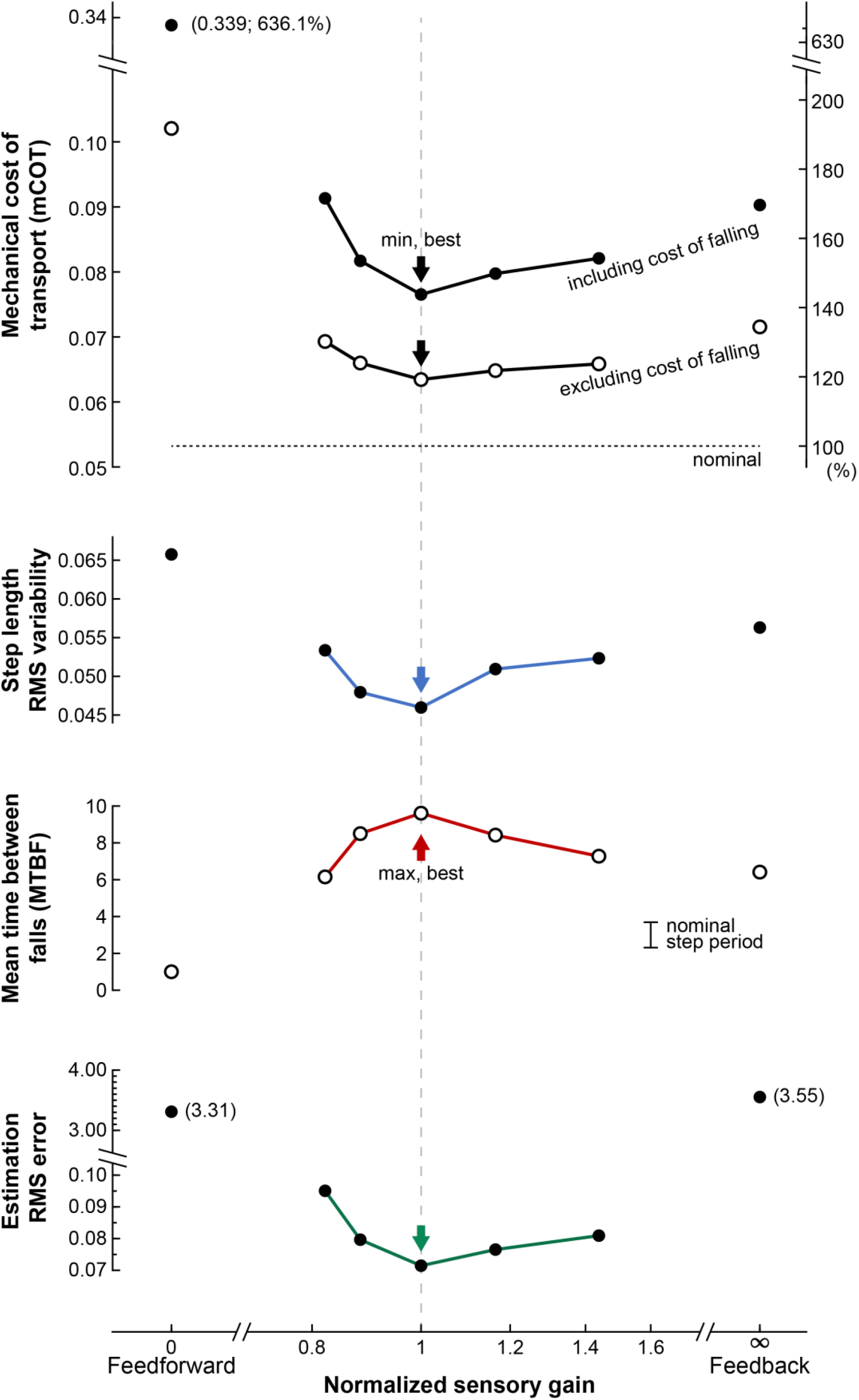
Walking performance under noisy conditions, as a function of sensory gain, in terms of mechanical cost of transport (mCOT), step length variability, mean time between falls (MBTF), and state estimator error. Normalized sensory gain varies between extremes of pure feedforward (to the left) and pure feedback (to the right), with 1 corresponding to theoretical optimum 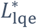. Formally, normalized gain is defined as 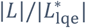, where | · | denotes matrix norm. Vertical arrow indicates best performance (minimum for all measures except maximum for MTBF). For all gains, model was simulated with a fixed combination of process and sensor noise as input to multiple trials, yielding ensemble average measures. Mechanical cost of transport (mCOT) was defined as positive work divided by body weight and distance travelled, and step variability as root-mean-square (RMS) variability of step length. Falling takes time and dissipates mechanical energy, and so mCOT was computed both including and excluding losses from falls (work, time, distance.

As expected of optimal estimation, best performance was found for the gain *L* equal to theoretically optimized 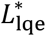 (Figure 5). We had designed 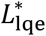 with linear quadratic estimation (LQE) based on the covariances of process and sensor noise. Using that gain in nonlinear simulation, the mechanical cost of transport was 0.077, somewhat higher than the nominal 0.053 without noise. Step length variability was 0.046 *l*, and the model experienced occasional falls, with MTBF (mean time between falls) of about 9.61 *g*^−0.5^*l*^0.5^ (or about 7.1 steps). This optimal case served as a basis for comparisons with other values for gain *L*.

Other values for sensory gain generally resulted in poorer performance (Figure 5). Over the range of gains examined (normalized sensory gain 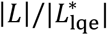 ranging 0.82 – 1.44), the performance measures worsened on the order of about 10%. This suggests that, in a noisy environment, a combination of feedforward and feedback is important for achieving precise and economical walking, and for avoiding falls. Moreover, the optimal combination for performance can be designed using control and estimation principles.

### Fictive locomotion emerges

Although the CPG model normally interacts with the body, it was also found to produce fictive locomotion even with peripheral feedback removed (Figure 6). Here we considered two types of biological sensors, referred to as “error feedback” and “measurement feedback” sensors. Error feedback refers to sensors that can distinguish unexpected perturbations from intended movements (44). For example, some muscle spindles and fish lateral lines (45) receive corollary efferent signals (e.g. gamma motor neurons in mammals, alpha in invertebrates (46)) that signify intended movements, and could be interpreted as effectively computing an error signal within the sensor itself (45). Measurement feedback sensors refers to those without efferent inputs (e.g., nociceptors, golgi tendon organs, cutaneous skin receptors, and other muscle spindles (47)), that provide information more directly related to body movement. Both types of sensors are considered important for locomotion, and so we examined the consequences of removing either type.

**Figure 6.**
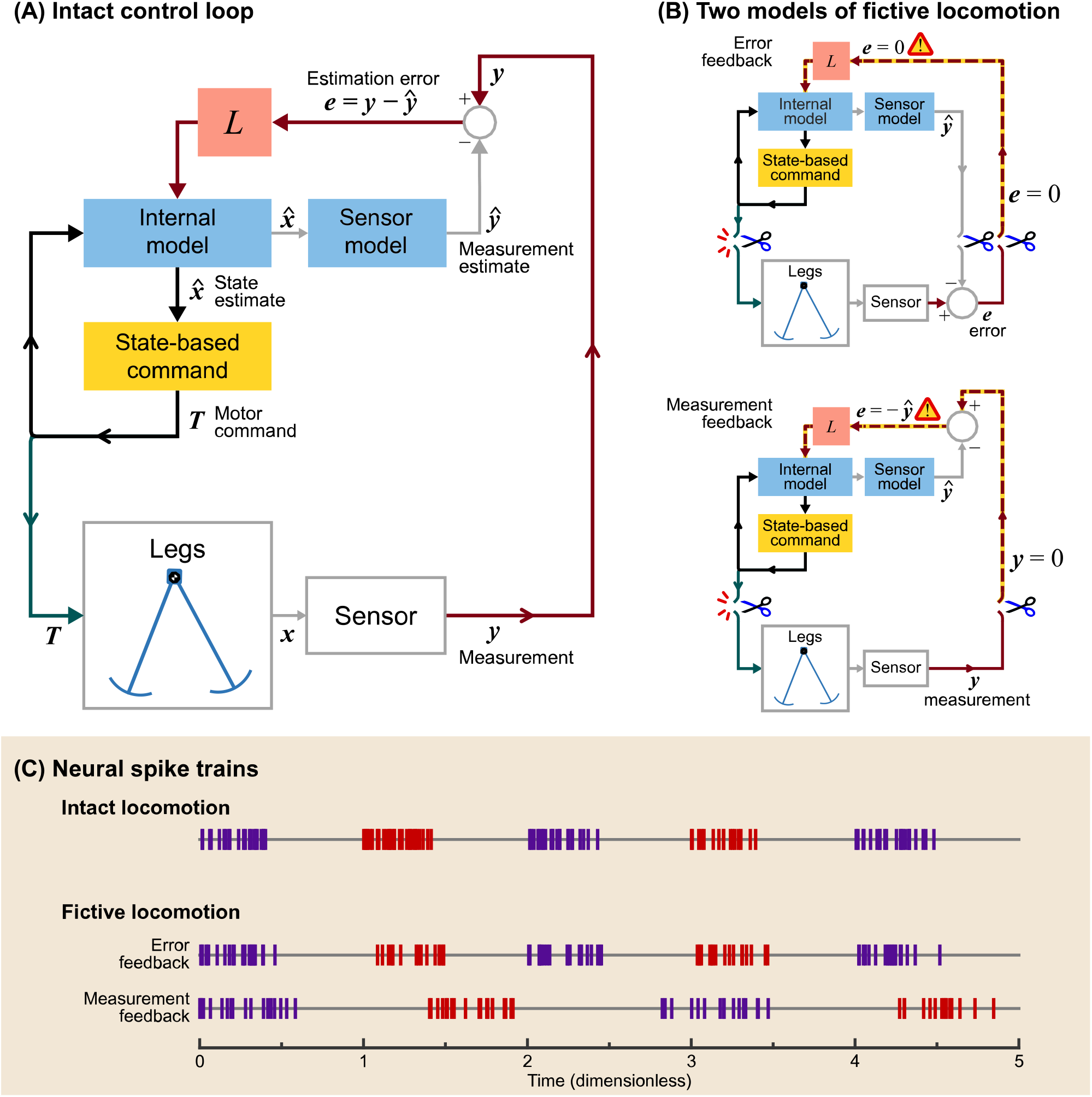
Emergence of fictive locomotion from CPG model. (A) Block diagram of intact control loop, where sensory measurements *y* and estimation error *e* are fed into internal model. Motor command *T* drives the legs and (through efference copy) the internal model of legs. (B) Two models of fictive locomotion, where sensory feedback is removed in two ways. Error feedback refers to sensors that receive efferent copy as inhibitory drive (e.g., some muscle spindles). Removal of error (dashed line) results in sustained fictive rhythm, due to feedback between internal model and state-based command. Measurement feedback refers to other, more direct sensors of limb state ***x***. Removal of such feedback can also produce sustained rhythm from internal model of legs and sensors, interacting with state-based command. (C) Simulated motor spike trains show how fictive locomotion can resemble intact. Measurement feedback case produces slower and weaker rhythm than error feedback.

These cases were modeled by disconnecting different components of the closed-loop system. This is best illustrated by redrawing the CPG (Figure 4) more explicitly as a traditional state estimator block diagram (Figure 6A). The case of removing error feedback (Figure 6B) was modeled by disconnecting error signal *e*, so that the estimator would run in an open-loop fashion, as if the state estimate were always correct. Despite this disconnection, there remained an internal loop between the estimator internal model and the state-based command generator, that could potentially sustain rhythmic oscillations. The case of removing measurement feedback (Figure 6C) was modeled by disconnecting afferent signal *y*, and reducing estimator gain by about half, as a crude representation of highly disturbed conditions. There remained an internal loop, also potentially capable of sustained oscillations. We tested whether either case would yield a sustained fictive rhythm, illustrated by transforming the motor command *T* into neural firing rates using a Poisson process.

We found that removal of both types of sensors still yielded sustained neural oscillations (Figure 6C), equivalent to fictive locomotion. In the case of error feedback, the motor commands from the isolated CPG were equivalent to the intact case without noise in terms of frequency and amplitude. In the case of measurement feedback, simulations still produced periodic oscillations, albeit with slower frequency and reduced amplitude compared to intact. The state estimator tended to drive estimate 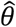 toward zero, and how this altered state estimation affected the final motor commands was quite dependent on the specifics of the state-based motor command.

## Discussion

We have examined how central pattern generators could optimally integrate sensory information to control locomotion. Our CPG model offers an adjustable gain on sensory feedback, to allow for continuous adjustment between pure feedback control to pure feedforward control, all with the same nominal gait under perfect conditions. The model is compatible with previous neural oscillator models, while also being designed through optimal state estimation principles. Simulations reveal how sensory feedback becomes critical under noisy conditions, although not to the exclusion of intrinsic, neural dynamics. In fact, a combination of feedforward and feedback is generally favorable, and the optimal combination can be designed through standard estimation principles. Estimation principles apply quite broadly, and could be readily applied to other models, including ones far more complex than examined here. The state estimation approach also suggests new interpretations for the role of CPGs in animal or robot locomotion.

One of our most basic findings was that the extremes of pure feedforward or pure feedback control each performed relatively poorly in the presence of noise (Figure 3). Pure feedforward control, driven solely by an open-loop rhythm, was highly susceptible to falling as a result of process noise. The general problem with a feedforward or time-based rhythm is that a noisy environment can disturb the legs from their nominal motion, so that the nominal command pattern is mismatched for the perturbed state. Under noisy conditions, it is better to trigger motor commands based on feedback of actual limb state, rather than time. But feedback also has its weaknesses, in that noisy sensory information can lead to noisy commands. The solution is to combine both feedforward and feedback together, modulated by sensory feedback gain *L*. A more uncertain environment (higher process noise) would favor higher gain, and noisier sensors would favor reduced gain. And for a given combination of noise, the theoretically optimal gain would be expected to minimize estimation error, and in turn yield best gait performance (normalized sensory gain = 1 in Figure 5). This is expected because theoretically, optimal control also typically calls for optimal state estimation (28,48). And empirically, imprecise visual information can induce variability in foot placement (49) and poorer walking economy (50). As expected, the present model walks best with the optimal trade-off between noise effects.

Our model also explains how neural oscillators can be interpreted as state estimators (Figure 4). Previous CPG oscillator models have incorporated sensory feedback for locomotion (15–22), but have not generally defined the feedback gain for optimal performance, based on mechanistic principles to combine feedforward and feedback. We have re-interpreted neural oscillator circuits in terms of state estimation (Figure 4), and shown how the gain can be determined in a principled manner, to minimize estimation error (Figure 5). The nervous system has long been interpreted in terms of internal models, for example in central motor planning and control (51–53) and in peripheral sensors (44). Here we apply internal model concepts to CPGs, for better locomotion performance.

This interpretation also explains fictive locomotion as an emergent behavior. We observed persistent CPG activity despite removal of sensors (and either error or measurement feedback; Figure 6), but this was not because the CPG was in any way intended to produce rhythmic timing. Rather, fictive locomotion was a side effect of a state-based motor command, in an internal feedback loop with a state estimator, resulting in an apparently time-based rhythm (Figure 6B). Others have cautioned that CPGs should not be interpreted as generating decisive timing cues (54–56), especially given the critical role of peripheral feedback in timing (9,57,58). In normal locomotion, central circuits and periphery act together in a feedback loop, and so neither can be assigned primacy. The present model operationalizes this interaction, demonstrates its optimality for performance, and shows how it can yield both normal and fictive locomotion.

This study argues that it is better to control with state rather than time. The kinematics and muscle forces of locomotion might appear to be time-based trajectories driven by an internal clock. But another view is that the body and legs comprise a dynamical system dependent on state (described e.g. by phase-plane diagrams, Figure 2), such that the motor command should also be a function of state. That control could be continuous as examined here, or include discrete transitions between circuits (e.g., (8)), and could be optimized or adapted through a variety of approaches, such as optimal control (28), dynamic programming (27), iterative linear quadratic regulators (59), and deep reinforcement learning (26)). But with noisy sensors (30,31), state-based control typically also requires state estimation (30,60), which introduces intrinsic dynamics and the possibility of sustained internal oscillations. Thus, robots with state-driven optimal control and estimation might also exhibit CPG-like fictive behavior, despite having no explicit time-dependent controls.

State estimation may also be applicable to movements other than locomotion. The same circuitry employed here (Figure 4) could easily contribute a state estimate 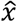 for any state-dependent movements, whether rhythmic (35), non-rhythmic, or discrete. In our view, persistent oscillations could be the outcome of state estimation with an appropriate state-based command for the *α* motoneuron (see Methods). But the same half-center circuitry could be active and contribute to other movements that use non-locomotory, state-based commands. It is certainly possible that biological CPGs are indeed specialized purely for locomotion alone, but the state estimation interpretation suggests the possibility of a more general, and perhaps previously unrecognized, role in other movements.

The present optimization approach may offer insight on neural adaptation. Although we have explicitly designed a state estimator here, we would also expect a generic neural network, given an appropriate objective function, to be able to learn the equivalent of state estimation. The learning objective could be to minimize error of predicted sensory information, or simply locomotion performance such as cost of transport. Moreover, our results suggest that the eventual performance and control behavior should ultimately depend on body dynamics and noise. A neural system adapting to relatively low process noise (and high sensor noise) would be expected to learn and rely heavily on an internal model. Conversely, relatively high process noise (and low sensor noise) would rely more heavily on sensory feedback. A limitation of our model is that it places few constraints on neural representation, because there are many ways (or “state realizations” (48)) to achieve the same input-output function for estimation. But the importance and effects of noise on adaptation are hypotheses that might be testable with artificial neural networks or animal preparations.

There are, however, cases where state estimation may not apply. State estimation applies best to systems with inertial dynamics or momentum. Examples include inverted pendulum gaits with limited inherent (or passive dynamic; (42)) stability and pendulum-like leg motions (35). The perturbation sensitivity of such dynamics makes state estimation more critical. But other organisms and models may have well-damped limb dynamics and inherently stable body postures, and thus benefit less from state estimation. There may also be task requirements that call for fast reactions with short synaptic delays, or organismal, energetic, or developmental considerations that limit the complexity of neural circuitry. Such concerns might call for reduced-order internal models (48), or even their elimination altogether, in favor of faster and simpler pure feedforward or feedback. A more holistic view would balance the principled benefits of internal models and state estimation against the practicality, complexity, and organismal costs.

There are a number of limitations to this study. The “Anthropomorphic” walking model does not capture three-dimensional motion and multiple degrees of freedom in real animals. We used such a simple model because it is unlikely to have hidden features that could produce the same results for unexpected reasons. We also modeled extremely simple sensors, without representing the complexities of actual biological sensors. The estimator also used a constant, linear gain, and could be improved with nonlinear estimator variants. We also used a particularly simple, state-based command law, which was designed more for robustness than for economy. Better economy could be achieved by powering gait with precisely-triggered, trailing-leg push-off (35), rather than the simple hip torque applied here. However, the timing is so critical that feedforward conditions (low sensory gains, Figure 5) would fall too frequently to yield meaningful economy or step variability measures. We therefore elected for more robust control to allow a range of feedforward through feedback to be compared (Figure 5). But even with more economical control, we would still expect optimal performance to correspond with optimal gain.

Our principal contribution has been to reconcile optimal control and estimation with biological CPGs. Evidence of fictive locomotion has long shown that neural oscillators produce timing and amplitude cues. But pre-determined timing is also problematic for optimal control in unpredictable situations (55), leading some to question why CPG oscillators should dictate timing (54,56). To our knowledge, previous CPG models have not included process or sensor noise in control design. Such noise is simply a reality of non-uniform environments and imperfect sensors. But it also yields an objective criterion for uniquely defining control and estimation parameters. The resulting neural circuits resemble previous oscillator models and can produce and explain nominal, noisy, or fictive locomotion. In our interpretation, there is no issue of primacy between CPG oscillators and sensory feedback, because they interact optimally to deal with a noisy world.

## Method

Details of the model and testing are as follows. The CPG model is first described in terms of neural, half-center circuitry, which is then paired with a walking model with pendulum-like leg dynamics. The walking gait is produced by a state-based command generator, which governs how state information is used to drive motor neurons. The model is subjected to process and sensor noise, which tend to cause the gait to be imprecise and subject to falling. The CPG is then re-interpreted as an optimal state estimator, for which sensory feedback gain and internal model parameters may be designed, as a function of noise characteristics. The model is then simulated over multiple trials to computationally evaluate its walking performance as a function of sensory gain. It is also simulated without sensory feedback, to test whether it produces fictive locomotion.

### CPG architecture based on Matsuoka oscillator

The CPG consists of two, mutually-inhibiting half-center oscillators, receiving a tonic descending input (Figure 4A). Each half-center has second-order dynamics, described by states *u*_*i*_ and *v*_*i*_. This is equivalent to a primary Matsuoka neuron with states for a membrane potential and adaptation or fatigue (14). Locomotion requires relatively longer time constants than is realistic for a single biological neuron, and so each model neuron here should be regarded as shorthand for a network of biological neurons with adaptable time constants and synaptic weights, that in aggregate produce first- or second-order dynamics of appropriate time scale. The state *u*_*i*_ produces an output *q*_*i*_ that can be fed to other neurons. In addition, we included two types of auxiliary neurons (for a total of three neurons per half-center): one for accepting the ground contact input (*c*_*i*_, with value 1 when in ground contact and 0 otherwise for leg *i*), and the other to act as an alpha (*α*_*i*_) motoneuron to drive the leg. We used a single motoneuron to generate both positive and negative (extensor and flexor) hip torques, as a simplifying alternative to including separate rectifying motoneurons.

Each half-center receives a descending command and two types of sensory feedback. The descending command is a tonic input *s*, which determines the walking speed. Sensory input from the corresponding leg includes continuous and discrete information. The continuous feedback contains information about leg angle from muscle spindles and other proprioceptors (61), which could be modeled as leg angle *y*_*i*_ for measurement feedback, or error *e*_*i*_ for error feedback sensors. The discrete information is about ground contact *c*_*i*_ sent from cutaneous afferents (62).

The primary neuron’s second-order dynamics are described by two states. The state *u*_*i*_ is mainly affected by its own adaptation, a mutually inhibiting connection from other neurons, sensory input, and efference copy of the motor commands. The second state *v*_*i*_ has a decay term, and is driven by the same neuron’s *u*_*i*_ as well as sensory input. This is described by the following equations, inspired by (14) and previous robot controllers designed for rhythmic arm movements (e.g., (63) and walking (22)):

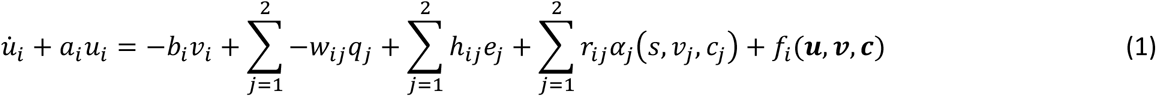

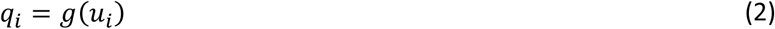

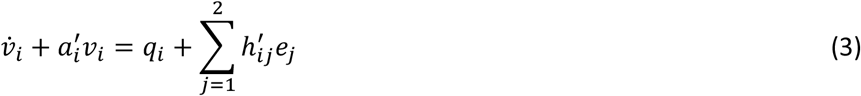

where there are several synaptic weightings: decays *a*_*i*_ and 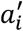, adaptation gain *b*_*i*_, mutual inhibition strength *w*_*ij*_ (weighting of neuron *i*’s input from neuron *j*’s output, where *w*_*ii*_ = *0*), an output function *g*(*u*_*i*_) (set to identity here), sensory input gains *h*_*ij*_ and 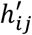, and efference copy strength *r*_*ij*_. The neuron also receives efference copy of its associated motor command *α*_*j*_(*s, v*_*j*_, *c*_*j*_), which depends on neuron state, descending drive and ground contact. There are also secondary, higher-order influences summarized by the function *f*_*i*_ (***u, v, c***), which have a relatively small effect on neural dynamics but are part of the state estimator. The network parameters for such CPG oscillators are traditionally set through a combination of design rules of thumb and hand-tuning, but here nearly all of the parameters here will be determined from an optimal state estimator, as described below.

### Walking model with pendulum dynamics

The system being controlled is a simple bipedal model walking in the sagittal plane (Fig. 3A). The passive dynamics of pendulum-like legs (42) are actively actuated by added torque inputs (the “Anthropomorphic Model,” (41)), and energy is dissipated mainly with the collision of leg with ground in the step-to-step transition. The dissipation determines the amount of positive work required each step. In humans, muscles perform much of that work, which in turn accounts for much of the energetic cost of walking (39).

The walking model is described mathematically as follows. The equations of motion may be written in terms of vector ***θ*** ≜ [*θ*_1_,*θ*_2_]^*T*^ as

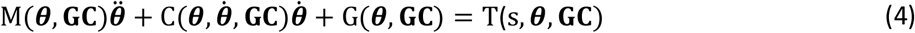

where M is the mass matrix, C describes centripetal and Coriolis effects, G contains position-dependent moments such as from gravity, **GC**≜ [GC_1_,GC_2_]^*T*^ contains ground contact, and **T** ≜ [*T*_1_,*T*_2_]^*T*^ contains hip torques exerted on the legs (State-based control, below). The equations of motion depend on ground contact because each leg alternates between stance and swing leg behaviors, inverted pendulum and hanging pendulum, respectively. We define each matrix to switch the order of elements at heel-strikes, so that equation of motion can be expressed in the same form.

At heelstrike, the model experiences a collision with ground affecting the angular velocities. This is modeled as a perfectly inelastic collision. Using impulse-momentum, the effect may be summarized as the linear transformation

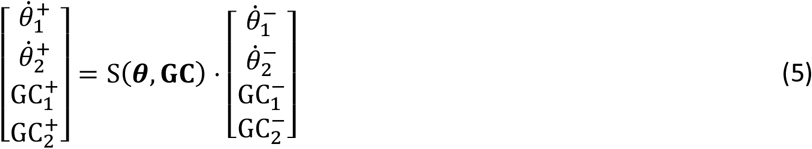

where the plus and minus signs (‘+’ and ‘–’) denote just after and before impact, respectively. The ground contact states are switched such that the previous stance leg becomes and swing leg, and vice versa.

The resulting gait has several characteristics relevant to the CPG. First, the legs have pendulum-like inertial dynamics, which allow much of the gait to occur passively. For example, each leg swings passively, and its collision with ground automatically induces the next step of a periodic cycle (42). Second, inertial dynamics integrate forces over time, such that disturbances can disrupt timing and cause falls. This sensitivity could be reduced with overdamped joints and low-level control, but humans are thought to have significant inertial dynamics (38). And third, inertial dynamics are retained in most alternative models with more degrees of freedom and more complexity (e.g., (24,26)). We believe most other dynamical models would also benefit from feedback control similar in concept to presented here.

### State-based motor command generator

The model produces state-dependent hip torque commands to the legs. Of the many ways to power a dynamic walking model (e.g., (41,64–66)), we apply a constant extensor hip torque against the stance leg, for its parametric simplicity and robustness to perturbations. The torque normally performs positive work (Fig. 3B) to make up for collision losses, and could be produced in reaction to a torso leaned forward (not modeled explicitly here; (42)). The swing leg experiences a hip torque proportional to swing leg angle (Fig. 3B, 3C), with the effect of tuning the swing frequency (35). This control scheme is actually suboptimal for economy, and is selected for its robustness. Optimal economy actually requires perfectly-timed, impulsive forces from the legs (40), and has poor robustness to the noisy conditions examined here. The present non-impulsive control is much more robust, and can still have its performance optimized by appropriate state estimation.

The overall torque command *T*_*i*_ for leg *i* is used as the motor command *α*_*i*_, and may be summarized as

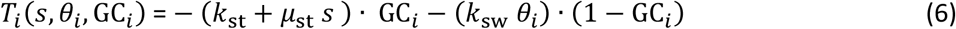

where the stance phase torque is increased from the initial value *k*_*st*_ by the amount proportional to the descending command *s* with gain *μ*_*st*_. The swing phase torque has gain *k*_*sw*_ for the proportionality to leg angle *θ*_*i*_.

There are also two higher level types of control acting on the system. One is to regulate walking speed, by slowly modulating the tonic, descending command *s* (Eqn. 6). An integral control is applied on *s*, so keep attain the same average walking speed despite noise, which would otherwise reduce average speed. The second type of high-level control is to restart the simulation after falling. When falling is detected (as a horizontal stance leg angle), the walking model is reset to its nominal initial condition, except advanced one nominal step length forward from the previous footfall location. No penalty is assessed for this re-set process, other than additional energy and time wasted in the fall itself. We quantify the susceptibility to falling with a mean time between failures (MTBF), and report overall energetic cost in two ways, including and excluding failed steps. The wasted energy of failed steps is ignored in the latter case, resulting in lower reported energy cost.

### Noise model with process and sensor noise

The walking dynamics are subject to two types of noisy disturbances, process and sensor noise (Figure 3). Both are modeled as zero-mean, Gaussian white noise. Process noise *n*_*x*_ (with covariance *N*_*x*_) acts as an unpredictable disturbance to the states, due to external perturbations or noisy motor commands. Sensor or measurement noise *n*_*y*_ (with covariance *N*_*y*_) models imperfect sensors, and acts additively to the sensory measurements *y*. The errors induced by both types of noise are unknown to the CNS controller, and so both tend to reduce performance.

The noise covariances were set so that the model would be significantly affected by both types of noise. We sought levels sufficient to cause significant risk of falling, so that good control would be necessary to avoid falling while also achieving good economy. Process noise was described by covariance matrix *N*_*x*_, with diagonals filled with variances of noisy accelerations, which had standard deviations of *0*.*0*1*5* (*g/l*) for stance leg, *0*.1*6* (*g/l*) for swing leg. Sensor noise covariance *N*_*y*_ was also set as a diagonal matrix with both entries of standard deviation *0*.1. Noise was implemented as a spline interpolation of discrete white noise sampled at frequency of 16 (*g/l*)^0.5^ (well above pendulum bandwidth) and truncated to no more than ±3 standard deviations.

### State estimator with internal model of dynamics

A state estimator is formed from an internal model of the leg dynamics being controlled (see block diagram in Fig. 4), to produce a prediction of the expected state 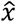 and sensory measurements *ŷ* (with the hat symbol ‘^’ denoting an internal model estimate). Although the actual state is unknown, the actual sensory feedback *y* is known, and the expectation error *e* = *y* − *ŷ* may be fed back to the internal model with negative feedback (gain *L*) to correct the state estimate. Estimation theory shows that regulating error *e* toward zero also tends to drive the state estimate towards actual state (assuming system observability, as is the case here; e.g., (48)). This may be formulated as an optimization problem, where gain *L* is selected to minimize the mean-square estimation error. Here we interpret the half-center oscillator network as such an optimal state estimator, the design of which will determine the network parameters.

The estimator equations may be described in state space. The estimator states are governed by the same equations of motion as the walking model (Eqns. 4, 5), with the addition of the feedback correction. Again using hat notation for state estimates, the nonlinear state estimate equations are

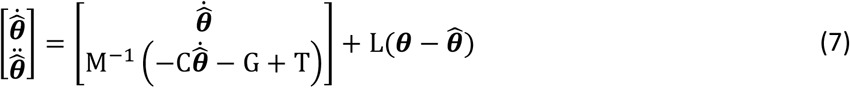

We used standard state estimator equations to determine a constant sensory feedback gain *L*. This was done by linearizing the dynamics about a nominal state, and then designing an optimal estimator based on process and sensor noise covariances (*N*_*x*_ and *N*_*y*_) using standard procedures (“lqe” command in Matlab, The MathWorks, Natick, MA). This yields a set of gains that minimize mean-square estimation error 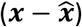, for an infinite horizon and linear dynamics. The constant gain was then applied to the nonlinear system in simulation, with the assumption that the resulting estimator would still be nearly optimal in behavior. Another sensory input to the system is ground contact GC_i_, a boolean variable. The state estimator ignores measured GC_i_ for pure feedforward control (zero feedback gain *L*), but for all other conditions (non-zero *L*), any sensed change in ground contact overrides the estimated ground contact 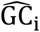. When the estimated ground contact state changes, the estimated angular velocities are updated according to the same collision dynamics as the walking model (eqn. 5 except with estimated variables).

The state estimate is applied to the state-based motor command (Eqn. 6). Although the walking control was designed for actual state information (*θ*_*i*_, GC_*i*_), for walking simulations it uses the state estimate instead:

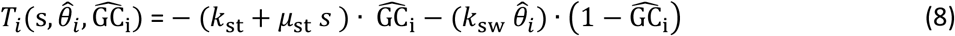

As with the estimator gain, this also requires an assumption. In the present nonlinear system, we assume that the state estimate may replace the state without ill effect, a proven fact only for linear systems (certainty-equivalence principle, (28,67)). Both assumptions, regarding gain *L* and use of state estimate, are tested in simulation below.

### Theoretical equivalence between neural oscillator and state estimator

Having fully described the walking model in terms of control systems principles, the equivalent half-center oscillator model may be determined (Figure 4B). The identical behavior is obtained by re- interpreting the neural states in terms of the dynamic walking model states,

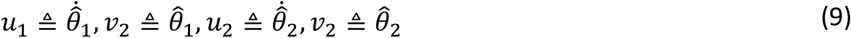

along with the neural output function defined as identity,

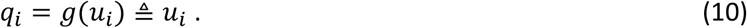

In addition, motor command and ground contact state are defined to match state-based variables (Eqn. 6)

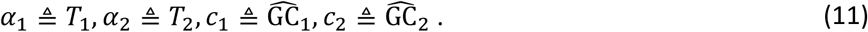

The synaptic weights and higher-order functions (Eqns. 1 – 3) are defined according to the internal model equations of motion (Eqn. 7),

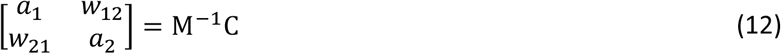

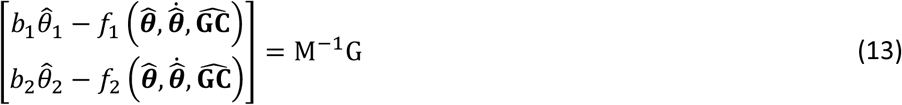

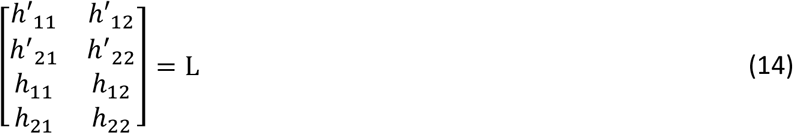

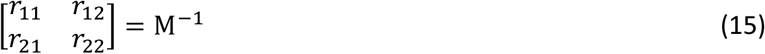

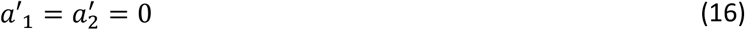

Because the mass matrix and other variables are state dependent, the weightings above are state dependent as well. The functions *f*_1_ and *f*_2_ are higher-order terms, which could be considered optional; omitting them would effectively yield a reduced-order estimator.

The result of these definitions is that the half-center neuron equations (Eqns. 1 – 3) may be rewritten in terms of 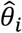 and 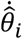, to illustrate how the network models the leg dynamics and receives inputs from sensory feedback and efference copy:

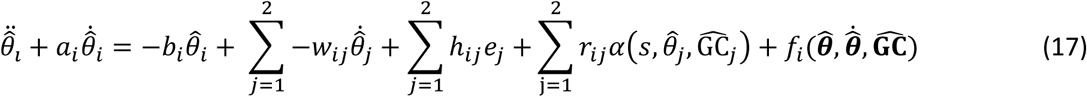

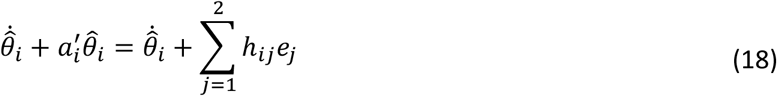

The above may be interpreted as an internal model of the stance and swing leg as pendulums, with pendulum phasing modulated by error feedback *e*_*j*_ and efference copy of the motor command (plus small nonlinearities due to inertial coupling of the two pendulums).

### Parametric effect of varying sensory feedback gain *L*

The sensory feedback gain is selected using state estimation theory, according to the amount of process noise and sensor noise. High process noise, or uncertainty about the dynamics and environment, favors a higher feedback gain, whereas high sensor noise favors a lower feedback gain. The ratio between the noise levels determines the optimal linear quadratic estimator gain 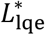 (Matlab function “lqe”). A constant gain was determined based on a linear approximation for the leg dynamics, an infinite horizon for estimation, and a stationarity assumption for noise (48). In simulation, the state estimator was implemented with nonlinear dynamics, assuming this would yield near-optimal performance.

It is thus instructive to evaluate walking performance for a range of feedback gains. Setting *L* too low or too high would be expected to yield poor performance. Setting *L* equal to the optimal LQE gain 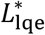 would be expected to yield approximately the least estimation error, and therefore the most precise control (e.g. (68)). In terms of gait, more precise control would be expected to reduce step variability and mechanical work, both of which are related to metabolic energy expenditure in humans (e.g., (50)). The walking model is also prone to falling when disturbed by noise, and optimal state estimation would be expected to reduce the frequency of falling.

We performed a series of walking simulations to test the effect of varying the feedback gain. The model was tested with 20 trials of 100 steps each, subjected to pseudorandom process and sensor noise of fixed covariance (*W* and *V*, respectively). In each trial, walking performance was assessed with mechanical cost of transport (mCOT, defined as positive mechanical work per body weight and distance travelled; e.g., (43)), step length variability, and mean time between falls (MTBF) as a measure of walking robustness (also referred to as Mean First Passage Time (69)). The sensory feedback gain 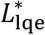 was first designed in accordance with the experimental noise parameters, and then the corresponding walking performance was evaluated. Additional trials were performed, varying sensory feedback gain *L* with lower and higher than optimal values to test for a possible performance penalty. These sub-optimal gains were determined by re-designing the estimator with sensor noise *ρV* (*ρ* between 10^-4 and 10^0.8, with smaller values tending toward pure feedforward and larger toward pure feedback). This procedure guarantees stable closed-loop estimator dynamics, which would not be the case if the matrix 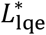 were simply scaled higher or lower. For all trials, the redesigned *L* was tested in simulations using the fixed process and sensor noise levels. The overall *sensory gain* was quantified with a scalar, defined as the L2 norm (largest singular value) of matrix *L*, normalized by the L2 norm of 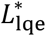.

We expected that optimal performance in simulation would be achieved with gain *L* close to the theoretically optimal LQE gain, 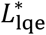. With too low a gain (*L* = *0*, feedforward Figure 1A), the model would perform poorly due to sensitivity to process noise, and with too a high gain (*L* → ∞, feedback Figure 1C), it would perform poorly due to sensor noise. And for intermediate gains, we expected performance to have an approximately convex bowl shape, centered about a minimum at or near 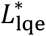. These differences were expected from noise alone, as the model was designed to yield the same nominal gait regardless of gain *L*. Simulations were necessary to test the model, because its nonlinearities do not admit analytical calculation of performance statistics.

### Evaluation of fictive locomotion

We tested whether the model would produce fictive locomotion with removal of sensory feedback. Disconnection of feedback in a closed-loop control system would normally be expected to eliminate any persistent oscillations. But estimator-based control actually contains two types of inner loops (Figure 6A), both of which could potentially allow for sustained oscillations in the absence of sensory feedback. However, the emergence of fictive locomotion and its characteristics depend on what kind of sensory signal is removed. We considered two broad classes of sensors, referred to producing error feedback and measurement feedback, with different expectations for the effects of their removal.

Some proprioceptors relevant to locomotion, including some muscle spindles and fish lateral lines (45), could be regarded as producing error feedback. They receive corollary discharge of motor commands, and appear to predict intended movements, so that the afferents are most sensitive to unexpected perturbations. The comparison between expected and actual sensory output largely occurs within the sensor itself, yielding error signal *e* (Figure 6B). Disconnecting the sensor would therefore disconnect error signal *e*, and would isolate an inner loop between state-based command and internal model. The motor command normally sustains rhythmic movement of the legs for locomotion, and would also be expected to sustain rhythmic oscillations within the internal model. Fictive locomotion in this case would be expected to resemble the nominal motor pattern.

Sensors that do not receive corollary discharge could be regarded as direct sensors, in that they relay measurement feedback related to state. In this case, disconnecting the sensor would be equivalent to removing measurement *y*. This isolates two inner loops, both the command-and-internal-model loop above, as well as a sensory prediction loop between sensor model and internal model. The interaction of these loops would be expected to yield a more complex response, highly dependent on parameter values. Nonetheless, we would expect that removal of *y* would substantially weaken the sensory input to the internal model, and generally result in a weaker or slower fictive rhythm.

We tested for the existence of sustained rhythms for both extremes of error feedback and measurement feedback. Of course, actual biological sensors within animals are vastly more diverse and complex than this model. But the existence of sustained oscillations in extreme cases would also indicate whether fictive locomotion would be possible with some combination of different sensors within these extremes.

## End Note

The source code for the simulation, supplementary table & video are available in a public repository at: https://bitbucket.org/hsRyu/cpg_biped_walker_ryu_kuo/src/master/

## Acknowledgements

This work was supported in part by NSERC (Discovery Award and CRC Tier I), Dr. Benno Nigg Research Chair, and University of Calgary BME Graduate Program.

## Reference

1. Brown TG. On the nature of the fundamental activity of the nervous centres; together with an analysis of the conditioning of rhythmic activity in progression, and a theory of the evolution of function in the nervous system. The Journal of Physiology. 1914 Mar 31;48(1):18–46.

2. Wilson DM. The Central Nervous Control of Flight in a Locust. Journal of Experimental Biology. 1961 Jun 1;38(2):471–90.

3. Wilson DM, Wyman RJ. Motor Output Patterns during Random and Rhythmic Stimulation of Locust Thoracic Ganglia. Biophys J. 1965 Mar;5(2):121–43.

4. Grillner S. Locomotion in vertebrates: central mechanisms and reflex interaction. Physiological Reviews. 1975 Apr 1;55(2):247–304.

5. Feldman AG, Orlovsky GN. Activity of interneurons mediating reciprocal 1a inhibition during locomotion. Brain Research. 1975 Feb 7;84(2):181–94.

6. Sherrington CS. Flexion-reflex of the limb, crossed extension-reflex, and reflex stepping and standing. The Journal of Physiology. 1910 Apr 26;40(1–2):28–121.

7. Pringle JWS. The Reflex Mechanism of the Insect Leg. Journal of Experimental Biology. 1940 Jan 1;17(1):8–17.

8. Büschges A. Sensory control and organization of neural networks mediating coordination of multisegmental organs for locomotion. J Neurophysiol. 2005 Mar;93(3):1127–35.

9. Bässler U, Büschges A. Pattern generation for stick insect walking movements--multisensory control of a locomotor program. Brain Res Brain Res Rev. 1998 Jun;27(1):65–88.

10. Liu CJ, Fan Z, Seo K, Tan XB, Goodman ED. Synthesis of Matsuoka-Based Neuron Oscillator Models in Locomotion Control of Robots. In: 2012 Third Global Congress on Intelligent Systems. 2012. p. 342–7.

11. Habib MK, Keigo Watanabe, Kiyotaka Izumi. Biped locomotion using CPG with sensory interaction. In: 2009 IEEE International Symposium on Industrial Electronics. 2009. p. 1452–7.

12. Auddy S, Magg S, Wermter S. Hierarchical Control for Bipedal Locomotion using Central Pattern Generators and Neural Networks. In: 2019 Joint IEEE 9th International Conference on Development and Learning and Epigenetic Robotics (ICDL-EpiRob). 2019. p. 13–8.

13. Cristiano J, García MA, Puig D. Deterministic phase resetting with predefined response time for CPG networks based on Matsuoka’s oscillator. Robotics and Autonomous Systems. 2015 Dec 1;74:88–96.

14. Matsuoka K. Mechanisms of frequency and pattern control in the neural rhythm generators. Biol Cybernetics. 1987 Jul 1;56(5–6):345–53.

15. Iwasaki T, Zheng M. Sensory Feedback Mechanism Underlying Entrainment of Central Pattern Generator to Mechanical Resonance. Biol Cybern. 2006 Apr 1;94(4):245–61.

16. Nassour J, Hénaff P, Benouezdou F, Cheng G. Multi-layered multi-pattern CPG for adaptive locomotion of humanoid robots. Biol Cybern. 2014 Jun 1;108(3):291–303.

17. Tsuchiya K, Aoi S, Tsujita K. Locomotion control of a biped locomotion robot using nonlinear oscillators. In: Proceedings 2003 IEEE/RSJ International Conference on Intelligent Robots and Systems (IROS 2003) (Cat No03CH37453) [Internet]. Las Vegas, NV, USA: IEEE; 2003 [cited 2019 Apr 30]. p. 1745–50. Available from: http://ieeexplore.ieee.org/document/1248896/

18. Morimoto J, and, Hyon S, Cheng G, Bentivegna D, Atkeson CG. Modulation of simple sinusoidal patterns by a coupled oscillator model for biped walking. In: Proceedings 2006 IEEE International Conference on Robotics and Automation, 2006 ICRA 2006. 2006. p. 1579–84.

19. Kimura H, Fukuoka Y, Cohen AH. Adaptive Dynamic Walking of a Quadruped Robot on Irregular Terrain Based on Biological Concepts. :16.

20. Righetti L, Ijspeert AJ. Pattern generators with sensory feedback for the control of quadruped locomotion. In: 2008 IEEE International Conference on Robotics and Automation [Internet]. Pasadena, CA, USA: IEEE; 2008 [cited 2019 Apr 30]. p. 819–24. Available from: http://ieeexplore.ieee.org/document/4543306/

21. Bliss T, Iwasaki T, Bart-Smith H. Central Pattern Generator Control of a Tensegrity Swimmer. IEEE/ASME Trans Mechatron. 2013 Apr;18(2):586–97.

22. Endo G, Morimoto J, Nakanishi J, Cheng G. An empirical exploration of a neural oscillator for biped locomotion control. In: IEEE International Conference on Robotics and Automation, 2004 Proceedings ICRA ‘04 2004. 2004. p. 3036-3042 Vol.3.

23. Alexander RM. Optima for Animals [Internet]. Princeton, NJ: Princeton University Press; 1996 [cited 2020 Sep 5]. Available from: https://press.princeton.edu/books/paperback/9780691027982/optima-for-animals

24. Geyer H, Herr H. A Muscle-Reflex Model That Encodes Principles of Legged Mechanics Produces Human Walking Dynamics and Muscle Activities. IEEE Transactions on Neural Systems and Rehabilitation Engineering. 2010 Jun;18(3):263–73.

25. Heess N, TB D, Sriram S, Lemmon J, Merel J, Wayne G, et al. Emergence of Locomotion Behaviours in Rich Environments. arXiv:170702286 [cs] [Internet]. 2017 Jul 7 [cited 2018 Feb 16]; Available from: http://arxiv.org/abs/1707.02286

26. Peng XB, Berseth G, Yin K, Van De Panne M. DeepLoco: Dynamic Locomotion Skills Using Hierarchical Deep Reinforcement Learning. ACM Trans Graph. 2017 Jul;36(4):41:1–41:13.

27. Bellman R. The theory of dynamic programming. Bull Amer Math Soc. 1954 Nov;60(6):503–15.

28. Bryson AE, Ho Y-C, Siouris GM. Applied Optimal Control: Optimization, Estimation, and Control. IEEE Transactions on Systems, Man, and Cybernetics. 1979;6(9):366–7.

29. Bryson AE. Applied Optimal Control: Optimization, Estimation and Control. CRC Press; 1975. 500 p.

30. Kuindersma S, Deits R, Fallon M, Valenzuela A, Dai H, Permenter F, et al. Optimization-based locomotion planning, estimation, and control design for the atlas humanoid robot. Autonomous Robots. 2016;40(3):429–455.

31. Wooden D, Malchano M, Blankespoor K, Howardy A, Rizzi AA, Raibert M. Autonomous navigation for BigDog. In: 2010 IEEE International Conference on Robotics and Automation. 2010.. 4736–41.

32. Kuo AD. An optimal state estimation model of sensory integration in human postural balance. J Neural Eng. 2005 Sep;2(3):S235–249.

33. Kuo AD. An optimal control model for analyzing human postural balance. IEEE Trans Biomed Eng. 1995 Jan;42(1):87–101.

34. Todorov E. Stochastic Optimal Control and Estimation Methods Adapted to the Noise Characteristics of the Sensorimotor System. Neural Computation. 2005 May 1;17(5):1084–108.

35. Kuo AD. The relative roles of feedforward and feedback in the control of rhythmic movements. Motor Control. 2002 Apr;6(2):129–45.

36. O’Connor SM. The relative roles of dynamics and control in bipedal locomotion. [Ann Arbor, MI]: University of Michigan; 2009.

37. Kuo AD, Donelan JM, Ruina A. Energetic consequences of walking like an inverted pendulum: step-to-step transitions. Exerc Sport Sci Rev. 2005;33:88–97.

38. Alexander RM. Simple models of human motion. Applied Mechanics Reviews. 1995;48:461–9.

39. Donelan JM, Kram R, Kuo AD. Mechanical work for step-to-step transitions is a major determinant of the metabolic cost of human walking. Journal of Experimental Biology. 2002;205:3717–27.

40. Kuo AD. Energetics of actively powered locomotion using the simplest walking model. Journal of Biomechanical Engineering. 2002;124:113–20.

41. Kuo AD. A Simple Model of Bipedal Walking Predicts the Preferred Speed--Step Length Relationship. J Biomech Eng. 2001 Jun;123(3):264–9.

42. McGeer T. Passive dynamic walking. The International Journal of Robotics Research. 1990;9(2):62.

43. Collins S, Ruina A, Tedrake R, Wisse M. Efficient bipedal robots based on passive-dynamic walkers. Science. 2005 Feb 18;307(5712):1082–5.

44. Dimitriou M, Edin BB. Human muscle spindles act as forward sensory models. Current Biology. 2010;20(19):1763–7.

45. Straka H, Simmers J, Chagnaud BP. A New Perspective on Predictive Motor Signaling. Current Biology. 2018 Mar 5;28(5):R232–43.

46. Delcomyn F. Reflexes and pattern generation, Ch. 16. In: Foundations of Neurobiology. New York: W. H. Freeman; 1998. p. 383–400.

47. Iggo A. Handbook of sensory physiology. Volume II. Somatosensory system. 1973;851pp.

48. Kailath T. Linear Systems. Prentice-Hall; 1980. 712 p.

49. O’Connor SM, Kuo AD. Direction-dependent control of balance during walking and standing. J Neurophysiol. 2009 Sep;102(3):1411–9.

50. O’Connor SM, Xu HZ, Kuo AD. Energetic cost of walking with increased step variability. Gait & Posture. 2012 Mar 27;36(1):102–7.

51. Hwang EJ, Shadmehr R. Internal Models of Limb Dynamics and the Encoding of Limb State. J Neural Eng. 2005 Sep;2(3):S266–78.

52. Kawato M. Internal models for motor control and trajectory planning. Current Opinion in Neurobiology. 1999 Dec 1;9(6):718–27.

53. Uno Y, Kawato M, Suzuki R. Formation and control of optimal trajectory in human multijoint arm movement. Biol Cybern. 1989 Jun 1;61(2):89–101.

54. Bässler U. On the definition of central pattern generator and its sensory control. Biol Cybern. 1986 May 1;54(1):65–9.

55. Cruse H. The functional sense of central oscillations in walking. Biol Cybern. 2002 Apr;86(4):271– 80.

56. Pearson KG. Central Pattern Generation: A Concept Under Scrutiny. In: McLennan H, Ledsome JR, McIntosh CHS, Jones DR, editors. Advances in Physiological Research [Internet]. Boston, MA: Springer US; 1987 [cited 2019 Dec 4]. p. 167–85. Available from: https://doi.org/10.1007/978-1-4615-9492-5_10

57. Donelan JM, Pearson KG. Contribution of sensory feedback to ongoing ankle extensor activity during the stance phase of walking. Canadian journal of physiology and pharmacology. 2004;82(8–9):589–598.

58. Pearson KG. Proprioceptive regulation of locomotion. Current opinion in neurobiology. 1995;5(6):786–791.

59. Li W, Todorov E. Iterative linear quadratic regulator design for nonlinear biological movement systems. In: ICINCO (1). 2004. p. 222–229.

60. Barfoot TD. State Estimation for Robotics. Cambridge University Press; 2017. 381 p.

61. Proske U, Gandevia SC. The Proprioceptive Senses: Their Roles in Signaling Body Shape, Body Position and Movement, and Muscle Force. Physiological Reviews. 2012 Oct 1;92(4):1651–97.

62. Trulsson M. Mechanoreceptive afferents in the human sural nerve. Experimental Brain Research. 2001 Mar 5;137(1):111–6.

63. Williamson MM. Neural control of rhythmic arm movements. Neural Networks. 1998 Oct 1;11(7):1379–94.

64. Srinivasan M, Ruina A. Computer optimization of a minimal biped model discovers walking and running. Nature. 2005;439(7072):72–75.

65. Westervelt ER, Grizzle JW, Koditschek DE. Hybrid zero dynamics of planar biped walkers. IEEE Transactions on Automatic Control. 2003 Jan;48(1):42–56.

66. Spong MW. Passivity based control of the compass gait biped. IFAC Proceedings Volumes. 1999 Jul 1;32(2):506–10.

67. Simon HA. Dynamic Programming Under Uncertainty with a Quadratic Criterion Function. Econometrica (pre-1986); Evanston. 1956 Jan;24(1):74.

68. Rebula JR, Ojeda LV, Adamczyk PG, Kuo AD. The stabilizing properties of foot yaw in human walking. J Biomech. 2017 28;53:1–8.

69. Byl K, Tedrake R. Metastable Walking Machines. The International Journal of Robotics Research. 2009 Aug 1;28(8):1040–64.

